# Tracing Sticky Trails: The Historical Biogeography of Australia’s Glandular Goose-foots (*Dysphania*, Chenopodioideae, Amaranthaceae)

**DOI:** 10.64898/2026.06.01.728730

**Authors:** Anže Žerdoner Čalasan, Francesco Susca, Karol Krak, Bohumil Mandák, Gudrun Kadereit

## Abstract

**Aim:** The overarching aim of this study is to reconstruct the spatiotemporal evolutionary history of Australian *Dysphania*, including testing the littoral connection hypothesis, assessing the role of reticulation, and identifying the major drivers of diversification within the lineage.

**Location:** Australia and New Zealand

**Taxon:** *Dysphania*, Chenopodioideae, Amaranthaceae, Caryophyllales, Angiosperms

**Methods:** Using a DNA sequence dataset based on a target enrichment approach with custom baits designed for Chenopodioideae, we compared alternative biogeographic and ancestral habitat models to infer the spatiotemporal evolutionary history of Australian *Dysphania*. In addition, we applied several complementary analyses to assess concordance and conflict within our phylogenomic datasets, estimate ploidy levels, and compare ecological niches among closely related species.

**Results and Main conclusions:** Our results reveal a close evolutionary relationship between Sub-Saharan African and Australian desert ephemerals and indicate that Australian *Dysphania* originated through an ancestral reticulation event. The last common ancestor reached northwestern Australia during the Miocene, occupied riverine desert habitats, and migrated eastward with their expansion, potentially undergoing ecological speciation. Four major Australian clades subsequently diversified across Miocene to Pleistocene landscapes, from riverine deserts to salt lake mosaics, with divergence likely driven by salinity gradients, flood regimes, and microhabitat partitioning rather than polyploidisation or geographic isolation.

## Introduction

Contemporary Australian desert flora is thought to have evolved from coastal ancestors that were pre-adapted to saline environments. These adaptations allowed them to migrate into increasingly arid inland regions in the first place, where they survived on the shores of continuously receding palaeorivers that stepwise transformed into inland salt lakes during the Neogene due to increasing aridification. This process promoted the in-situ adaptations and facilitated speciation and range expansion events of these arid floral elements in absence of the competition from their temperate counterparts (Beadle, 1981; Burbidge, 1960; Crisp et al., 1999; Diels, 1906; Heads, 2013; Shmida, 1985). This ‘littoral connection hypothesis’ has been around for almost 100 years, but the next-generation sequencing data on desert elements needed to robustly test it, has been lacking. While major progress has been made in the past couple of years due to the development of new NGS techniques and improved sampling (Anderson et al., 2019; Cardillo et al., 2017; Cauz_JSantos et al., 2024; Hammer et al., 2021; Hancock et al., 2018; Hühn et al., 2024; Schmidt-Lebuhn & Smith, 2016), evolutionary histories of many prominent Australian desert elements such as Aizoaceae (*Gunniopsis*), Caesalpinioideae (*Acacia* and *Senna*), *Eremophila*, *Eucalyptus*, Frankeniaceae and Chenopodioideae, are still missing. One such lineage is the genus *Dysphania*, which represents an important but still poorly understood component of the Australian arid flora.

With only about 50 known species, *Dysphania* R.Br. is a relatively small genus that fits well morphologically and genetically within the subfamily Chenopodioideae (Fuentes-Bazan et al., 2012; Kadereit et al., 2010; Uotila et al., 2021). However, despite the evident similarities in habitus, pollen morphology and floral structure, *Dysphania* can be distinguished from other members of this subfamily by differences in pericarp indumentum and clear presence of multicellular glandular white hairs and/or yellow to orange subsessile glands (Sukhorukov & Zhang, 2013) that give this genus a characteristic aromatic odour that may persist in herbarium specimens for years (Uotila et al., 2021).

The genus *Dysphania* comprises almost exclusively annual species that are found on all continents with the exception of Antarctica, albeit, in similar light-rich, disturbed habitats (Figure 1). In Asia, *Dysphania* inhabits disturbed sites, well-drained sandy soils, grasslands at higher elevations and dry riverbed gravels (Sukhorukov et al., 2019). In North America, it occurs in riverbeds, dry lake beds, wastelands, along forest paths, sandy shores, and rocky soils or in light montane pine forests (Clemants & Mosyakin, 2004; Flora of North America; http://beta.floranorthamerica.org), and in South America, it favours arid or semi-arid regions, sandy soils, and ruderal areas near watercourses (Brignone, 2020; Flora de Cono Sur; http://conosur.floraargentina.edu.ar). In Africa, it occurs in montane areas, disturbed sites, and cattle-trampled soils (Sukhorukov et al., 2016). With 17 native species, the highest species diversity of this genus is found in Australia, where it thrives in semi-arid and arid regions on well-drained sandy soils, rocky outcrops, river banks, mudflats, seasonally flooded areas and a variety of soil types including skeletal, loamy and gypsum or calcareous soils, often around salt lakes and waterholes (Uotila et al., 2021; Wilson, 1984; Figure 1). Majority of *Dysphania* species confined to the arid zone occur on gypsum-rich saline soils (Symon, 2007). Surprisingly, despite evidently surviving in these rather inhospitable habitats, the ecological and physiological adaptations of *Dysphania* to these environments remain poorly understood. As annuals, the majority of these species tend to occur in great numbers after rainfall (McKenzie et al., 2007; Michael et al., 2010) and germination studies have supported these observations by showing that certain species germinate within hours after presence of water and produce long-lasting seed banks (Golos et al., 2016; Jurado & Westoby, 1992; Navie et al., 1996). In contrast to its closely related genus *Chenopodium*, whose main mode of evolution is allopolyploidisation (Krak et al., 2016; Mandák et al., 2018, 2026), speciation in *Dysphania* appears to be driven by different intrinsic mechanisms, as the species-rich Australian clade comprises only diploids. Furthermore, these different evolutionary modes are also reflected in their basic chromosome numbers: most American species have a base chromosome number of x = 8, whereas most diploid Eurasian and Australian species have x = 9 (Uotila et al., 2021).

**Figure 1:**
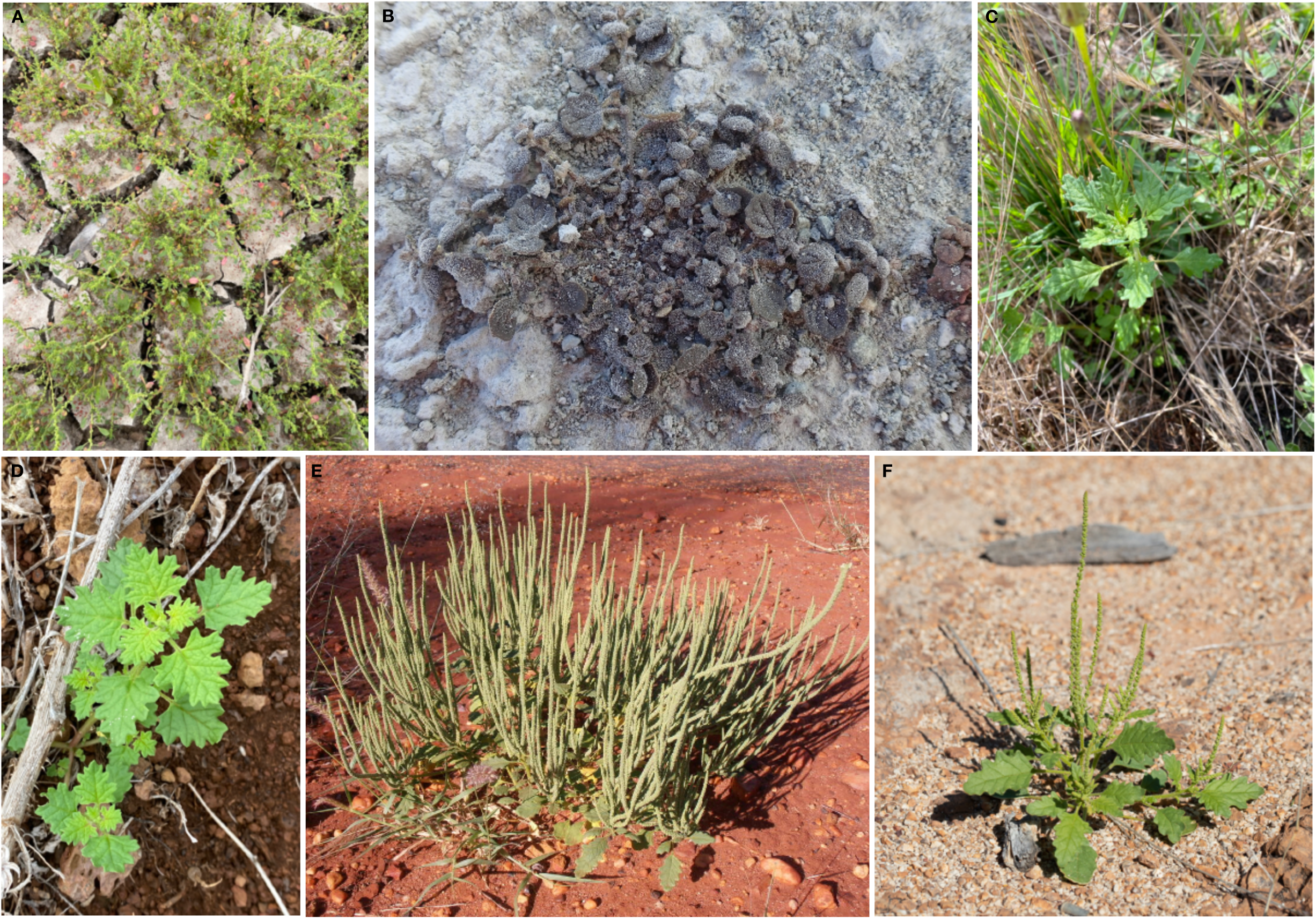
Diversity of Australian *Dysphania*: A, *D. glomulifera* occurring in high population numbers in an often flooded muddy area close to the Murray River (photo by Dr. Manfred Jusaitis, March 2023; location: –34.15, 140.35); B, *D. pusilla* from a dry, muddy area along a lake margin at high altitude in the New Zealand Alps (photo by Dr. Melissa Hutchison, January 2022; location omitted as this is a threatened taxon); C, widely distributed Australian native *D. pumilio* growing in an open patch of grass in northern New Zealand (photo by Joseph Knight, January 2025; location: –34.95, 173.37); D, widely distributed Australian native *D. carinata* growing along a hiking path on Maui Island (photo by John Starmer, October 2023; location: 20.64, –156.45); E, Australian native desert-dwelling *D. kalpari* on red sandy soil in full flower (photo by Dr. Tony Bean, April 2012; location: –25.96, 146.66); F, Australian native desert-dwelling *D. rhadinostachya* on sandy soil in full flower (photo by Dr. Kym Nicolson, June 2022; location: –22.56, 143.00).

With the exception of morphologically unusual *D. saxatile*, all Australian native taxa belong to the *Dysphania* sect. *Dysphania*. A recent phylogenetic analysis revealed that this section is monophyletic, whereas *D. saxatile* represents a single Australian species that clusters with South and East African representatives (Uotila et al., 2021). Phylogenetic relationships within Australian *Dysphania* largely corresponds to morphology, and the clades inferred from recent phylogenetic analyses (Uotila et al., 2021) can be readily distinguished by differences in perianth segment number and inflorescence architecture (Mosyakin, 2021). However, the temporal origin, biogeographic history, and evolutionary processes underlying diversification of Australian *Dysphania* remain largely unresolved. The main aims of this study are: i) to place the evolutionary history of Australian *Dysphania* in a spatiotemporal framework, ii) to test the littoral connection hypothesis against alternative inland or independent diversification scenarios in this taxon, iii) to test for reticulation events that may have given rise to *D. saxatilis* and iv) to investigate the drivers of diversification in Australian *Dysphania*.

## Materials and Methods

### Sampling

Here, we focused on the Australian and New Zealand taxa. Most of the material had already been analysed previously using Sanger sequencing by (Uotila et al., 2021), with specimens collected from the AD, BR, BRI, H, K, MJG, PERTH, RM and WU herbaria (Thiers, 2025). For this study, we supplemented the sample set with three additional accessions of the previously unsampled *D. pusilla*, which is endemic to New Zealand (Figure 1B), from CHR. The total number of analysed samples was 27 including all 17 species currently known to exist in Australia and New Zealand. The outgroup sampling comprised a transcriptome dataset of closely and distantly related genera across the Caryophyllales, which was previously published by (Morales-Briones, Kadereit, et al., 2021). Detailed voucher information is available in Supplementary File S1.

### Bait design

Low-copy gene targets were selected using the Sondovac tool (Schmickl et al., 2016), employing genome-skimming data from *Chenopodium ficifolium* (Belyayev et al., 2020) and *C. ficifolium* transcriptomes (Štorchová et al., 2019). The resulting set of candidate genes was then compared to the reference genomes of *C. suecicum* and *C. pallidicaule* (Jarvis et al., 2017), which represent diverse lineages within the genus *Chenopodium*. For more information, see Mandák et al., 2018. Only single-copy candidates were kept as targets for probe design. DAICEL Arbor Biosciences (Ann Arbor, Michigan, USA) designed and manufactured a 20K bait set targeting 6,198 exons belonging to 1,752 genes under Cat. No. 300116.v5.

### DNA extraction, library preparation and sequencing

Total DNA was extracted from 5-20 mg of dried leaf-material using the standard DNeasy Plant Mini Kit (QIAGEN, Venlo, Netherlands) following the manufacturer’s instructions and eluted in 100 µl of AE buffer (10 mM Tris, 0.5 mM EDTA, pH=9.0). DNA quality and quantity were assessed using a 1.0% agarose gel stained with ethidium bromide and the Invitrogen Qubit® 4 Fluorometer (Thermo Fisher Scientific), respectively. Library preparation was performed according to the standard NEBNext Ultra II FS Library Prep Kit protocol (New England Biolabs, Ipswich, MA, USA), with targeted inserts of approximately 300 bp and half-volume reactions. Prior to library preparation, non- or poorly degraded DNA samples were sheared with an M220 focused-ultrasonicator (Covaris, Massachusetts, USA). Following initial fragmentation and end repair, adapter ligation and size selection using AMPure XP magnetic beads were performed, along with a clean-up of the adapter-ligated DNA fragments. All libraries were dual-indexed using NEBNext® Multiplex Oligos for Illumina (New England Biolabs, Massachusetts, USA, #E6444). This was followed by PCR enrichment with 10 cycles, after which the resulting dual-indexed fragments were cleaned up using AMPure XP magnetic beads. The quantity and quality of the resulting libraries were estimated using the Invitrogen Qubit® 4 Fluorometer (Thermo Fisher Scientific) and the TapeStation 4150 High Sensitivity D1000 (Agilent Technologies), respectively. Up to 24 samples of similar quality and quantity were pooled and hybridised using the customised *Chenopodium* bait kit at 65 °C overnight according to the myBaits ‘Hybridisation Capture for Targeted NGS’ standard protocol, version 5.00 (Daicel Arbor Biosciences, Michigan, USA). After the washing step, the enriched libraries were resuspended and amplified through eight to ten PCR cycles using NEB Ultra II Q5 polymerase. The libraries were then cleaned up one last time with AMPure XP magnetic beads. The final quantity and quality of the libraries were assessed using the Invitrogen Qubit® 4 Fluorometer (Thermo Fisher Scientific) and the TapeStation 4150 High Sensitivity D1000 (Agilent Technologies) before they were sent out for sequencing. Sequencing was performed on a NovaSeq 6000 (Illumina Inc., San Diego, USA) using v1.5 chemistry and 150 bp paired-end reads at Novogene aiming at 2.3 Gb/sample.

### Data cleaning, assembly, extraction and alignment

The raw reads were deduplicated using clumpify.py script from BBMap/BBTools (Bushnell, 2014). Adaptor trimming, as well as quality trimming and filtering, were performed using the option ‘captus_assembly clean’ integrated into the CAPTUS pipeline (Ortiz et al., 2023) by removing read regions with phred quality scores under 20 as well as complete reads with average minimum phred quality score under 20 using Falco algorithm (De Sena Brandine & Smith, 2021). The cleaned reads were assembled using MEGAHIT (Li et al., 2015) integrated in ‘captus_assembly assemble’ pipeline with a minimum contig depth set to three, strict pruning of low-coverage edges and minimum contig length set to 100 bp. Loci were blasted against the target file used to generate the *Chenopodium* baits and extracted from CAPTUS assemblies using BLAT (Kent, 2002) and Scipio (Keller et al., 2008) incorporated in the ‘captus_assembly extract’. Only hits with the minimum percentage of 60% similarity and 50% of the reference protein length were retained. Finally, nuclear coding sequences and complete gene sequences (exons + introns) were extracted via ‘captus_assembly extract’ with a maximum number of retrieved paralogues set to five. Using a modified Python script we merged these with other outgroup Caryophyllales data generated by (Morales-Briones, Kadereit, et al., 2021). For these samples, we used previously generated orthologous sequences from which we extracted loci using the function ‘captus_assembly extract’ with the same target file, retaining only hits with the 75% minimum identity and 50% minimum coverage to the reference protein. The merged locus matrices were aligned using MACSE v.2.08 (Ranwez et al., 2018).

### Orthologue detection and final tree inference

Paralogue detection was carried out following the workflows of (Yang & Smith, 2014) and (Morales-Briones, Kadereit, et al., 2021), using the scripts available at https://bitbucket.org/dfmoralesb/target_enrichment_orthology. To streamline downstream analyses, we first masked both mono- and paraphyletic tips corresponding to the same taxon, retaining only the tip with the highest number of unambiguous characters in the trimmed alignment. Next, spurious tips in the trees were removed with TreeShrink v.1.3.9 (Mai & Mirarab, 2018), while excluding outgroup taxa. We then pruned paralogues sequentially using a monophyletic outgroup strategy, progressively expanding the outgroup until all outgroup taxa were included, thereby producing one-to-one orthologs in the absence of duplications. Only locus trees containing data for at least 25% of taxa were retained. From these trees, new locus alignments were generated, realigned using MACSE v.2.08 (Ranwez et al., 2018), and subsequently trimmed by removing columns with less than 30% occupancy using pxclsq v.1.3 (Brown et al., 2017). Phylogenetic trees for each locus were inferred with IQ-TREE v.2.1.2 (Minh et al., 2020) under the conditions described above. The resulting locus trees were rerooted, trimmed again with TreeShrink, and analysed using ASTRAL v.5.7.8 (Zhang et al., 2025).

### Gene discordance, orthogroup mapping and species network analysis

Target enrichment datasets are usually designed to capture primarily single-copy orthologs and avoid paralogs (Yang & Smith, 2014). This may limit the applicability of our orthogroup mapping inference method to detect whole genome duplications, which was originally developed for transcriptomic data. The reduced presence of paralogous sequences may consequently constrain the detection of duplication events. However, in line with the recommendations of (Morales-Briones, Gehrke, et al., 2021), we thus applied stringent filtering, retaining only loci with ≥80% ingroup taxon coverage, in order to minimise the potential for false positives arising from missing data. To infer potential whole genome duplication events, we extracted and mapped the orthogroups onto the final species tree using customised Python scripts ‘extract_clades.py’ and ‘map_dups_mrca.py’ from https://bitbucket.org/blackrim/clustering (Yang et al., 2018) with the ‘minimum input taxa’ value set to 30 (80%). To reduce the computational intensity of the analyses, the following two tests were run on a reduced taxon sample that included only the species of *Dysphania*, with *Suckleya* acting as the outgroup. To assess gene tree discordance, we used PhyParts (Smith et al., 2015), including only branches with bootstrap support above 70%, and complemented this analysis with quartet sampling (Pease et al., 2018). For the latter we used the concatenated alignment and a rooted species tree. Quartet sampling was run using IQ-TREE v2.3.1 (Minh et al., 2020) with 1,000 replicate samplings per internal branch.

To investigate potential reticulate evolutionary relationships among taxa, species network inference was performed using PhyloNet v3.8.2 (Wen et al., 2018). We first reduced the set of cleaned orthologous gene trees to a representative subset of 10 ingroup taxa capturing the major clades, with *S. suckleyana* designated as the outgroup. A total of 594 gene trees comprising all 11 taxa were used, and branches with local posterior probability support below 0.75 were collapsed prior to analysis. Species networks were inferred under the maximum pseudo-likelihood framework, allowing for 0 to 7 reticulation events. Each analysis was repeated 10 times, and the best-supported network was selected based on the highest pseudo-likelihood score. Final networks were visualized in Julia using the PyPlot.jl package (https://github.com/JuliaPy/PyPlot.jl).

### Time divergence estimation

Molecular dating was conducted using BEAUTi and BEAST v2.7.7 (Bouckaert et al., 2019) under an uncorrelated relaxed clock model (Drummond et al., 2006). The stem age of Amaranthaceae relied on pollen fossils of the genus *Chenopodipollis* from the Hell Creek Formation in North Dakota, dated to the Maastrichtian (72.2–66 my; (Nichols & Traverse, 1971), whereas the crown age of the Caryophyllales outgroup was further constrained to 130 ± 5 my following (Zuntini et al., 2024). To reduce computational demands, the alignment was filtered using a gene-shopping strategy in SortaDate (Smith et al., 2018). Gene trees were selected based on their similarity to the species tree, clock-likeness, and tree length, resulting in a reduced dataset of 25 loci. Using this dataset, along with a starting tree from the multispecies coalescent analysis, a GTR+Γ substitution model, and a Birth–Death tree prior to account for potential extinctions, we ran five independent MCMC chains for 100 million generations, sampling every 20,000 generations. Effective sample sizes (ESS) were assessed in Tracer v1.7.1 (Rambaut et al., 2018). To avoid overparameterisation, the +I option was omitted (Drummond & Bouckaert, 2015). The first 10% of trees were discarded as burn-in in TreeAnnotator v1.10.4 (Suchard et al., 2018), and the maximum clade credibility tree, with median node heights, was visualised in FigTree v1.4.4 (http://tree.bio.ed.ac.uk/software/figtree/).

### Biogeographic reconstruction

The biogeographic reconstruction was carried out in the R environment using ‘BioGeoBEARS’ v1.1.2 package (Matzke, 2018) in two steps. Firstly, to estimate from which region *Dysphania* arrived in Australia, the distribution areas were divided into the following entities: area A (Australia and New Zealand), area B (Eurasia), area C (Africa), area D (North America), and area E (South America). Native distribution areas for the taxa were estimated using Plant of the World Online (POWO; https://powo.science.kew.org). Secondly, to infer the dispersal of *Dysphania* within Australia, we subdivided the continent into distinct geographic entities. Here, the distribution areas for the taxa were estimated using Atlas of Living Australia (ALA) platform (https://ala.org.au; Belbin et al., 2021), and categorised into 20 ABA phytogeographical subregions (Ebach et al., 2015; González-Orozco et al., 2014) and New Zealand. To reduce the computational complexity of the analysis, we limited the number of biogeographic subregions to 10 by merging smaller subregion clusters of the Australian continent into larger units, based on an arbitrary threshold set in the dendrogram, based on species turnover (Ebach et al., 2015). However, as this resulted in clustering of two non-adjacent regions, Eastern Queensland was treated as an independent region from the Great Sandy Desert Interzone and the Central Desert. This resulted in the following areas: area A (Hampton, Southwest Interzone and Southwestern), area B (Western Desert), area C (Eastern Desert), area D (Adelaide, Eyre Peninsula and Nullarbor) area E (Southeastern, Tasmania, Victoria), area F (Eastern Queensland), area G (Central Queensland), area H (Great Sandy Desert Interzone and the Central Desert), area I (New Zealand) and area J (outside of Oceania; Supplementary File S2). As no species occurs in Kimberley Plateau, Arnhem Land, Atherton Plateau and Cape York Peninsula, Northern Australia was excluded from the analysis. Furthermore, taxa were assigned to a particular subregion only, if at least 10% of its occurrence points were found in it. As the most widely distributed species occurred in five subregions, this was set as the maximum number of subregions a taxon could have occupied. Six models have been tested (BAYAREALIKE, BAYAREALIKE+J, DEC, DEC+J, DIVALIKE, DIVALIKE+J). and the best-fit model was chosen following the corrected Akaike information criterion (AICc) and Akaike weight (AICc_wt).

### Ancestral habitat evolution

Furthermore, we wanted to investigate the evolution of the Australian *Dysphania* landscape preferences. We thus assigned individual taxa to different Australian land types primarily following (McDonald, 2020). They designated areas based on national, state, and regional floras, published ecological and systematic surveys, details of herbarium specimen collections available through ALA, and the authors’ personal observations, relying heavily on Mabbutt (1988). We refined their assignments by plotting the occurrence points of individual taxa across the 89 geographically distinct IBRA bioregions based on common climate, geology, landform, native vegetation, and species information (Australian Government Department of the Environment and Energy, 2016: https://www.dcceew.gov.au/environment/land/nrs/science/ibra) and then exclude IBRA areas in which less than 10% of the occurrence points per taxon were found. Ultimately, the distribution of *Dysphania* was categorised into the following areas: coastal, mesic plains and ranges, riverine deserts, desert lakes, sandy deserts, desert uplands, shield plains and stony deserts. Australian desert clay plains and limestone plains are not occupied by *Dysphania* and were thus excluded from the coding. Ancestral character estimation was performed in R using the packages phytools (Revell, 2012) and ape (Paradis & Schliep, 2019). For each habitat type, three discrete-state models i. e. equal rates (ER), symmetric (SYM), and all rates different (ARD) were fitted using maximum likelihood and model selection was based on AIC. Node and tip probabilities of habitat states were extracted and visualised as pie charts on the phylogeny, with normalised probabilities. Cumulative figures showing all habitats simultaneously were generated to illustrate the combined evolutionary history of habitat occupancy across the phylogeny.

### Ecological niche modelling

Closely related *Dysphania* species may share similar ecological niches. To estimate their ecological niches, the R packages dismo (Hijmans et al., 2024) and ENMeval (Kass et al., 2021) were used. All 19 bioclimatic variables were extracted from the CHELSA database (Brun et al., 2022), and occurrence data for individual species were downloaded from the Atlas of Living Australia platform (http://www.ala.org.au). Environmental variables were used in their original units without scaling or transformation. To reduce the computational complexity of the analyses, the Australian extent was defined as xmin = 110, xmax = 155, ymin = −45, and ymax = −10, and the number of random background points was capped at 5000. Prior to the selection of bioclimatic variables, a collinearity check was performed using variance inflation factors, and highly correlated variables were removed.

Four MaxEnt feature types (L, LQ, H, and LQH) were tested with the maxnet algorithm (Phillips et al., 2017), and the “checkerboard” method was used for spatial partitioning in cross-validation. Model selection followed the AICc principle, and the final figures were generated using the R package ggplot2 (Wickham, 2016). Niche similarities were estimated using the R package ecospat (Di Cola et al., 2017). Four ecologically similar and closely related *Dysphania* species groups were tested: *D. carinata*, *D. cristata*, *D. melanocarpa*, *D. pumilio* and *D. truncata* from clade 1, *D. plantaginella*, *D. simulans* and *D. sphaerosperma* from clade 2, *D. kalpari* and *D. rhadinostachya* from clade 3, and *D. glandulosa*, *D. glomulifera* and *D. littoralis* from clade 4. Species *D. valida* and *D. congestiflora* were excluded from the analysis due to the limited number of occurrence records.

## Results

### Statistics

The number of output reads ranged from approximately 6 to 23 million, with adapter content varying from 0.25% to 25%. Contig lengths ranged from 145 bp to almost 63000 bp, with N50 values between 519 bp and 1040 bp. The mean contig depth varied from 46x to 160x. After filtering, the final number of recovered nuclear loci ranged from 956 to 1,128, with an average locus length of 1148 bp, an average number of 94,6 informative positions per locus and an average locus number of 1,07 copies.

### Topology

A total of 1,370 loci passed paralog filtering. Supplementary File S3 illustrates the topology of the multiple species coalescent-based tree generated from these loci, which is well supported, with most local posterior probability (LPP) values at the maximum of 1.00. The genus *Dysphania* was retrieved as monophyletic (1.00 LPP) within the monophyletic Chenopodieae, which also included representatives of the genera *Krascheninnikovia* Gueldenst., *Chenopodium* L., *Spinacia* L. (1.00 LPP), *Oxybasis* Kar. & Kir, *Proatriplex* (W.A.Weber) Stutz & G.L.Chu, *Stutzia* E.H.Zacharias, *Extriplex* E.H.Zacharias and *Atriplex* L. (1.00 LPP), with *Suckleya* A.Gray as its sister genus (1.00 LPP).

*Dysphania* consisted of four lineages: *Dysphania* sect. *Adenois* (Moq.) Mosyakin & Clemants (1.00 LPP), including New World *D. anthelmintica* (L.) Mosyakin & Clemants and *D. ambrosioides* (L.) Mosyakin & Clemants; *Dysphania* sect. *Botryoides* (C.A.Mey.) Mosyakin & Clemants (1.00 LPP), including Old World *D. botrys* (L.) Mosyakin & Clemants, *D. procera* (Hochst. ex Moq.) Mosyakin & Clemants and *D. geoffreyi* Sukhor.; *Dysphania* sect. *Margaritaria* (Brenan) G. Kadereit, Sukhor. & Uotila (1.00 LPP), including African *D. congolana* (Hauman) Mosyakin & Clemants, *D. pseudomultiflora* (Murr) Verloove & Lambinon and *D. schraderiana* (Schult.) Mosyakin & Clemants, and Australian *D. saxatilis* (Paul G. Wilson) Mosyakin & Clemants; and Australian *Dysphania* sect. *Dysphania* R.Br. (1.00 LPP). Within this section, four clades were recognised. Clade 1 (1.00 LPP) included New Zealand *D. pusilla* (Hook.f.) Mosyakin & Clemants, Australian *D. melanocarpa* (J.M. Black) Mosyakin & Clemants, *D. truncata* (Paul G. Wilson) Mosyakin & Clemants, and three Australian natives with world-wide distributions: *D. carinata* (R.Br.) Mosyakin & Clemants, *D. cristata* (F. Muell.) Mosyakin & Clemants, and *D. pumilio* (R.Br.) Mosyakin & Clemants. Clade 2 (1.00 LPP) included *D. congestiflora* S.J. Dillon & A.S. Markey, *D. plantaginella* F. Muell., *D. sphaerosperma* Paul G. Wilson, and *D. simulans* F. Muell. & Tate. Clade 3 (1.00 LPP) consisted of *D. kalpari* Paul G. Wilson and *D. rhadinostachya* (F. Muell.) A.J. Scott, whereas clade 4 (1.00 LPP) included *D. glandulosa* Paul G. Wilson, *D. littoralis* R.Br., *D. valida* Paul G. Wilson, *D. platycarpa* Paul G. Wilson, and *D. glomulifera* (Nees) Paul G. Wilson (Supplementary File S3).

### Phyparts and quartet sampling

*Dysphania* sect. *Adenois* showed high gene-tree concordance (with about 66% of concordant trees; Supplementary File S4) and QS levels (QC=0.95), with most quartets being informative (QI=0.97; Supplementary File S5). While there was some conflict, it was not strongly skewed toward any particular alternative (QD=0.44). *Dysphania* sect. *Botryoides* showed moderately high gene-tree concordance (with about 50% trees being concordant; Supplementary File S4) and QS levels (QC=0.79), with less quartets being informative (QI=0.67) and partially skewed alternative (QD=0.79; Supplementary File S5). Despite maximum LPP statistical support, the *D.* sect. *Margaritaria* showed low levels of gene tree concordance with only about 25% of gene-trees supporting the topology and only a low proportion of informative quartets (QI=0.21). The branch leading to *D.* sect. *Dysphania* was well supported with about two thirds of concordant gene-trees and QC=0.84, and moderately well-informed (QI=0.66) with some directional conflict (QD=0.87). The proportion of main topological alternatives was almost negligible along all branches of the phylogeny (Supplementary File S4).

Within the *Dysphania* sect. *Dysphania*, gene-tree discordance analysis revealed modest to moderate levels of conflict (Supplementary File S4). Although the relationships between the clades showed moderate discordance, they also exhibited high levels of uninformative data (approximately 50%), despite maximum LPP statistical support. Short branch lengths connecting the clades, together with high levels of uninformative data (0.18<QI<0.2) suggested that much of the underlying signal was weak and the direction of discordance was often strongly skewed (0.85<QD<0.98; Supplementary File S5). All four clades suffered from low quartet informativeness (0.21<QI<0.55), and while three of the four clades showed moderate QS support (QC=0.62–0.72), clade 2 exhibited markedly lower QS support (QC=0.18).

### Orthogroup mapping and species network analysis

Orthogroup mapping indicated that approximately 18% of the orthogroups associated with the *Dysphania* lineage showed evidence of duplication, and around 12% of the orthogroups exhibited duplication at the node corresponding to the New World *Dysphania* sect. *Adenois*. The remaining values were comparatively low, spanning from 0.3% to 1.5% (Supplementary File S6).

Species network analyses indicated that the optimal model included three reticulation events (data not shown), with inferred inheritance probabilities that were largely symmetric, ranging from approximately 0.45 to 0.55 (Figure 2). Australian *Dysphania* sect. *Dysphania* was inferred to have originated from a reticulation event, receiving ∼47% of its genetic contribution from an ancestral lineage closely related to *D.* sect. *Margaritaria*. In turn, *D.* sect. *Margaritaria* itself was associated with a reticulation event involving a lineage sister to *D. saxatilis*, contributing ∼45% of its genetic material. Finally, *D. saxatilis* was also recovered as a reticulate lineage, deriving ancestry from both a lineage sister to the remaining Australian taxa and the same ancestral lineage closely related to *D.* sect. *Margaritaria* (Figure 2).

**Figure 2:**
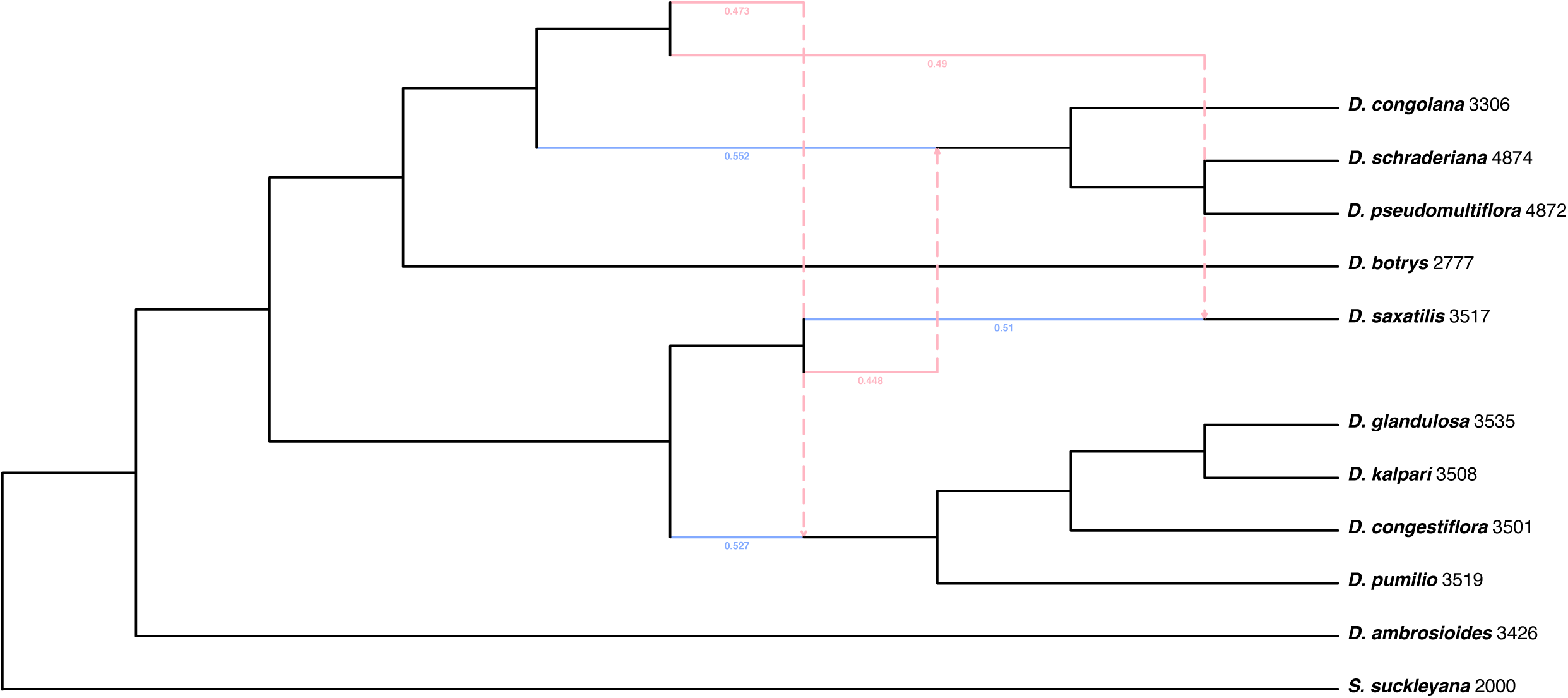
Phylogenetic network inferred under the MPL framework using a representative taxon set. Reticulation events shown in dark red, indicate secondary parental contributions, while those in blue represent primary parental contributions. Values on reticulation branches denote the estimated inheritance probabilities from each parental lineage.

### Time divergence estimation

The crown age of the Amaranthaceae family dated back to the Early Eocene, around 55 mya (48.1–60.9 mya, 95% HPD), whereas the crown age of the Chenopodioideae subfamily was estimated at approximately 35 mya (32.6–41.5 mya, 95% HPD), in the Late Eocene (Supplementary File S7). The last common ancestor of the genus *Dysphania* diverged at the beginning of the Miocene, around 23 mya (20.1–26.0 mya, 95% HPD), after which the genus began to diversify. The crown ages of the sections were all dated to the Late Miocene to Early Pleistocene: *Dysphania* sect. *Adenois*, 2.5 mya (1.9–3.1 mya, 95% HPD); *D.* sect. *Botryoides*, 6 mya (5.0–6.8 mya, 95% HPD); *D.* sect. *Margaritaria*, 7 mya (6.1–8.1 mya, 95% HPD); and *D.* sect. *Dysphania*, 5 mya (4.3–5.7 mya, 95% HPD; Supplementary File S7). The use of the uncorrelated relaxed clock model was justified, after assessing the coefficient of variation in all the trace files, which consistently exceeded 0.5. All effective sample size values exceeded 1000 pointing towards convergence (data not shown).

### Biogeographic reconstruction

On the continental scale, BioGeoBEARS v1.1.3 identified the DEC+J and DIVALIKE+J models as the best-fitting models (Supplementary File S8). According to these models, *Dysphania* arrived in Australia from North America, from where it subsequently spread to Eurasia and Africa (Figure 3). A statistically close second was the BAYAREALIKE+J model, which instead suggested that *Dysphania* reached Australia from Eurasia and then dispersed onward to Africa (Supplementary File S8; Figure 3).

**Figure 3:**
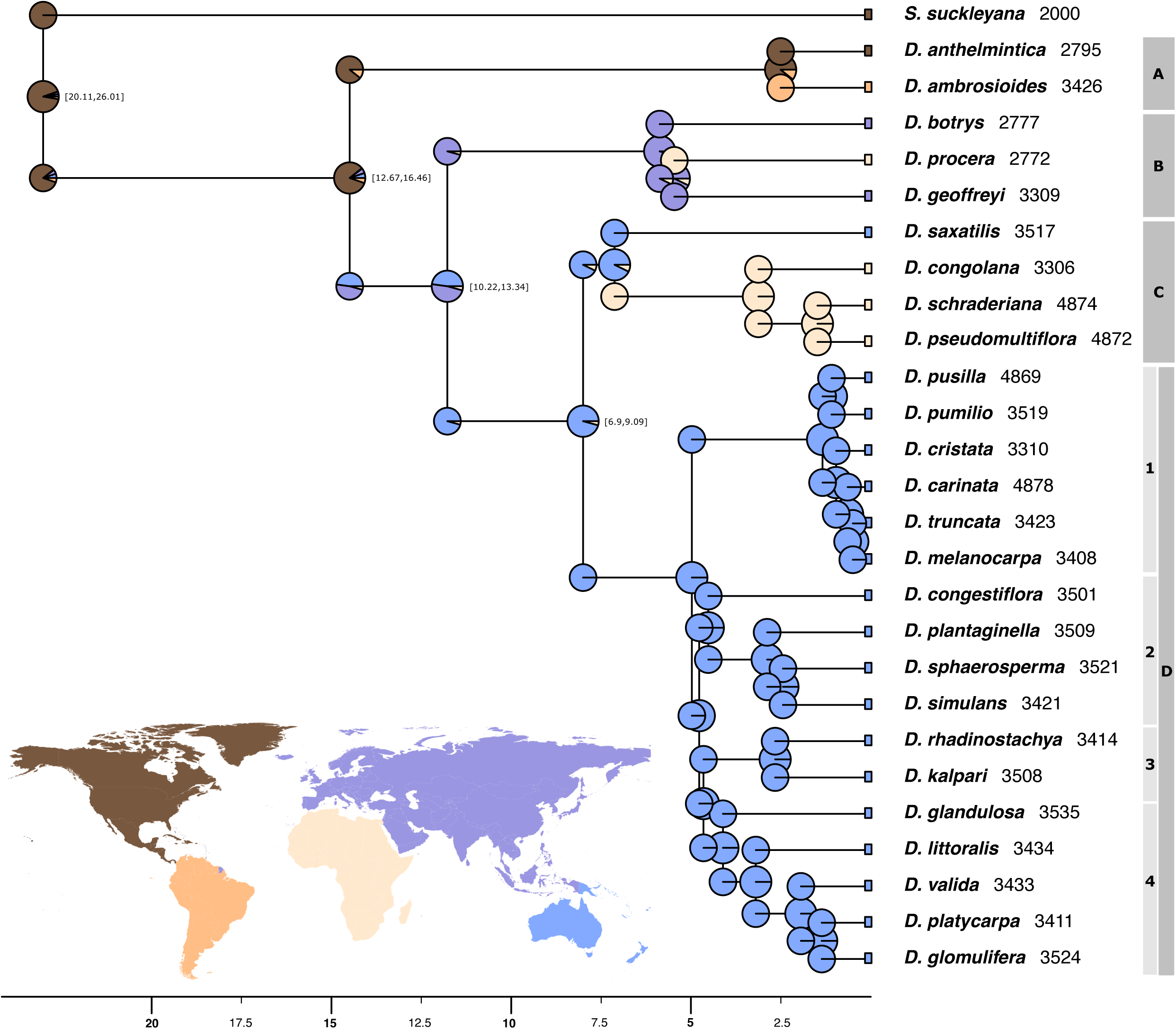
Spatiotemporal reconstruction of Australian *Dysphania* on the world scale. Columns next to the tip labels indicate individual *Dysphania* sections and clades within Australia. Abbreviations: A, *D.* sect. *Adenois*; B, *D.* sect. *Botrys*; C, *D.* sect. *Margaritaria*; D, *D.* sect. *Dysphania*. Pie charts show the inferred proportions of ancestral areas from a BioGeoBEARS analysis at the continental level, with colours corresponding to those on the world map in the lower left corner. The 95% HPD of some nodes is indicated in square brackets; ages for the remaining nodes are provided in Supplementary File S7. The x-axis is in millions of years.

At the Australian scale, the BAYAREALIKE+J model was recovered as the best fit (Supplementary File S8). According to this model, the last common ancestor of Australian *Dysphania* initially occupied the Western Desert (Figure 4A), from where it dispersed westward toward Africa (Figure 3) and eastward into the Eastern Desert (Figure 4B). From this eastern expansion, four additional range extension events were inferred dated to Pliocene and early Pleistocene: one into the coastal northeast region (Eastern Queensland; Figure 4C); two into the north-adjacent non-coastal Great Sandy Desert Interzone and the Central Desert, as well as into Central Queensland Figures 4D and 4E; and one into the coastal southwest regions (Hampton, Southwest Interzone, Southwestern; Figure 4F). Dispersal to the Mediterranean south, the temperate southeastern and eastern zones, and New Zealand took place later in the Pleistocene (Figure 4G).

**Figure 4:**
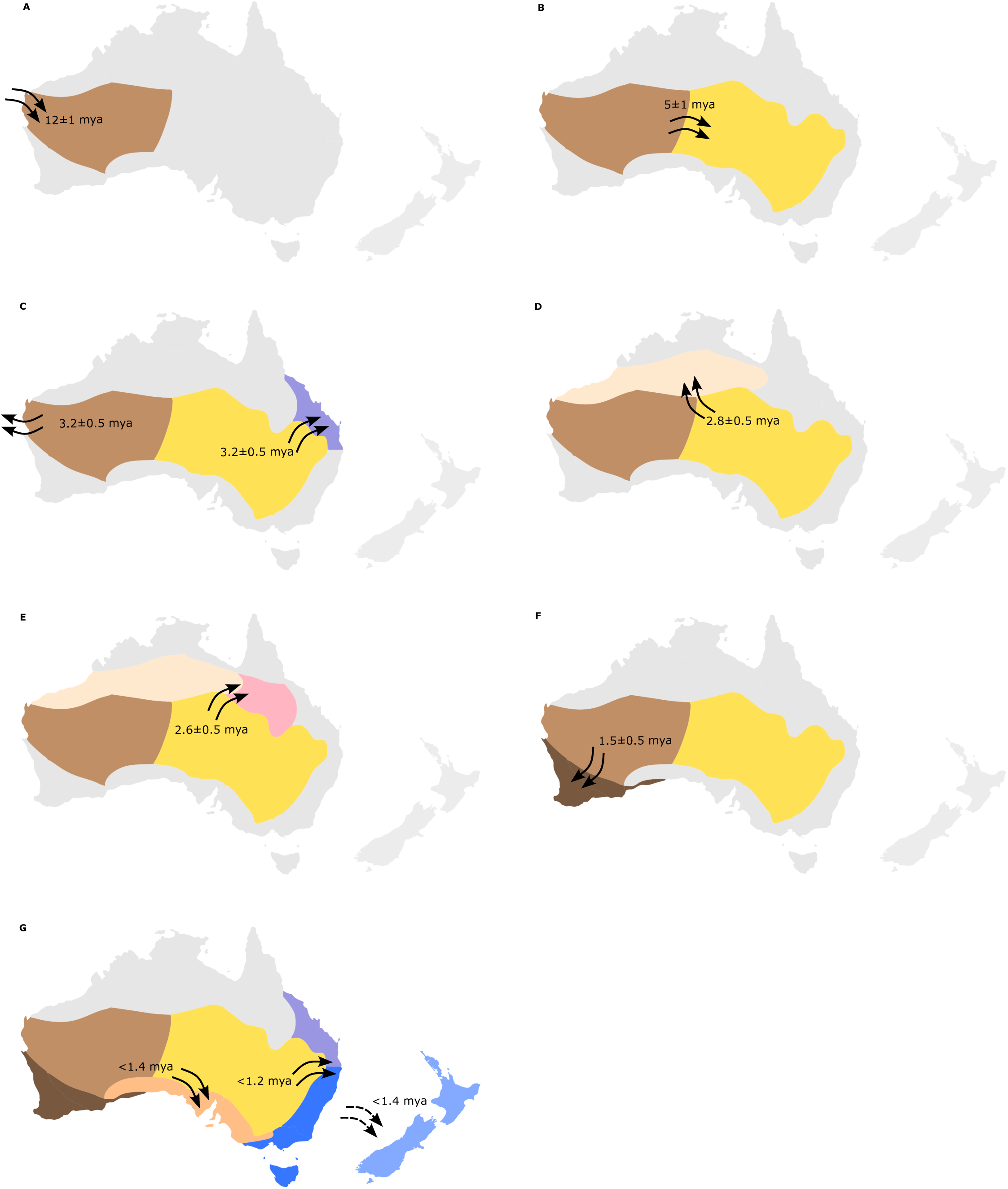
Spatiotemporal reconstruction of Australian *Dysphania* at the continental scale, inferred from a BioGeoBEARS analysis. Shaded areas indicate phytogeographical subregions according to Ebach et al. (2015), González-Orozco et al. (2014). Arrows show the directions of range expansions, with accompanying numbers representing the 95% HPD of each expansion as estimated from the divergence time analysis (Supplementary File S7).

### Landscape area reconstruction

The ARD model was selected as the best-fitting model for seven of the eight habitat states, with a minimal AIC difference of 1.88 compared to the SYM model for desert lake habitat (data not shown). Consistent with the two alternative models, the ancestral habitat of Australian *Dysphania* was reconstructed as riverine deserts, with >90% probability along the backbone, which remained one of the primary habitats for most species (Figure 5). One exception was clade 2, whose last common ancestor likely shifted its habitat preference to desert lakes, as indicated by high node probability (∼70%) in the ancestral reconstruction (Figure 5).

**Figure 5:**
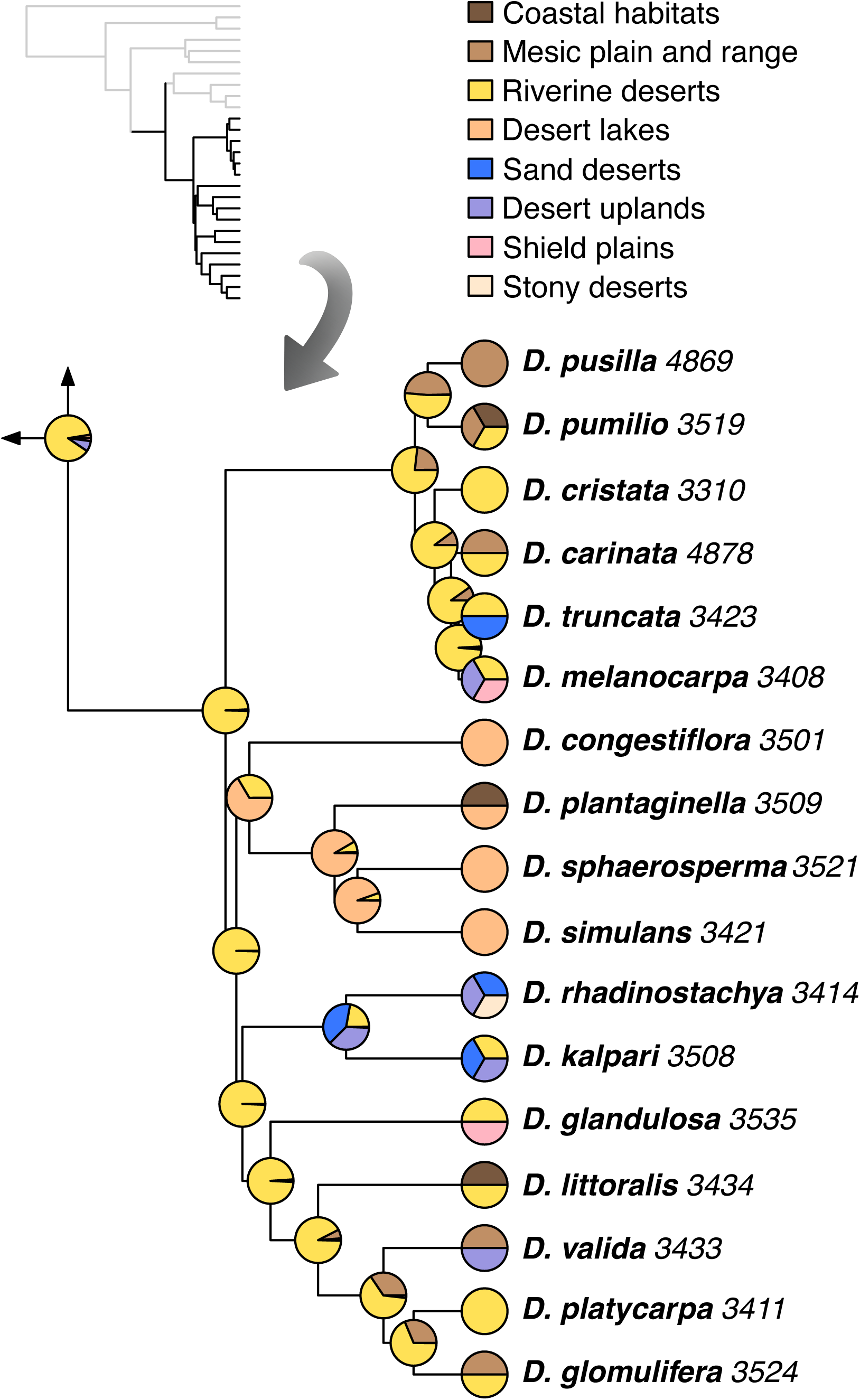
Ancestral habitat reconstruction of Australian *Dysphania* inferred under an ARD model. Pie charts at the tips indicate contemporary distributions, while pie charts at internal nodes show reconstructed ancestral habitats. Only the shaded portion of the tree in the upper left corner is displayed. Definitions of habitat types follow Mabbutt (1988) and McDonald (2020).

### Ecological niche modelling

The collinearity test revealed that, across all species, the least correlated variables were bio8, bio9, bio18, and bio19 (data not shown). Our analyses revealed distinct ecological patterns across *Dysphania* clades (Supplementary File S9). Within clade 1, *D. cristata* and *D. melanocarpa* shared the most similar environmental space (D = 0.67, I = 0.86), whereas *D. pumilio* occupied a largely distinct niche (D ≤ 0.008, I ≤ 0.064; (Supplementary File S10). *Dysphania truncata* overlapped moderately with *D. cristata* and *D. melanocarpa* but had minimal overlap with *D. carinata* and *D. pumilio*, and *D. carinata* showed a moderate to low overlap with the other species (D ≤ 0.245, I ≤ 0.44). In clade 2, *D. plantaginella* and *D. sphaerosperma* shared moderate to high environmental space (D = 0.62, I = 0.77), while *D. simulans* was largely distinct (D ≤ 0.002, I ≤ 0.04; (Supplementary File S10). In clade 3, the sister species *D. kalpari* and *D. rhadinostachya* exhibited moderate to high overlap (D = 0.56, I = 0.76). In clade 4, *D. glomulifera* and *D. littoralis* (D = 0.25, I = 0.41) showed the most similar, though still rather divergent, niches, whereas *D. platycarpa* was largely ecologically isolated (D ≤ 0.16, I ≤ 0.22; (Supplementary File S10).

## Discussion

### Justification of the tree topology

Three of the four morphologically and geographically well-defined sections exhibit high gene-tree concordance and concordance factors, indicating more or less coherent evolutionary lineages. The exception is *D.* sect. *Margaritaria*, which is neither morphologically nor geographically unified and shows low concordance (Supplementary Figure S4). This, combined with its hybrid origin inferred from our reticulation analyses (Figure 2), points to a complex evolutionary history of this clade. Although poorly known, this group comprises a single Australian species alongside African ones, rendering Australian *Dysphania* polyphyletic. The close relationship between Sub-Saharan African and Australian sclerophytes, xerophytes or desert ephemeral has been observed previously in many lineages, such as: *Atriplex* (Žerdoner Čalasan et al., 2022), Arctotidinae (McKenzie & Barker, 2008) and Gnaphalieae (Nie et al., 2016), Myrtaceae (Berger et al., 2016)*, Roepera* (Wu et al., 2018), *Salicornia* (Steffen et al., 2015), *Wahlenbergia* (Prebble et al., 2011)*, Tetragonia* (Klak et al., 2017) and *Trianthema* (Bohley et al., 2015).

As the sole Australian species in an otherwise African clade, *D. saxatilis* exhibits a combination of morphological traits: seed characters are reminiscent of clade 1 (formerly *Chenopodium* sect. *Orthosporum*), whereas less-lobed leaves, leafy inflorescences, four perianth segments, and vertical seeds align it with African *D. congolana*. This intermediate morphology is consistent with genomic evidence, which indicates that *D. saxatilis* originated through hybridisation between the ancestors of these two lineages (Figure 2). The morphology and ecology of species within the four clades constituting Australian *Dysphania* sect. *Dysphania* support their close evolutionary relationships. Species of clade 2 share spike-like inflorescences and predominantly grow along desert lakes or coastlines. By contrast, the sister species of clade 3 have four tepals and are predominantly found on poorly developed orthent soils. Clade 4 is ecologically more diverse, but all its species have flowers with reduced tepals that are packed tightly in axillary clusters. Although this section is of clear hybrid origin (Figure 2), this key morphological trait remains relatively stable across it, consistent with the hybrid signal being confined to the ancestral lineage rather than occurring between descendant species. It is tempting to speculate that this ancestral hybridisation event may have contributed to the relatively high number of taxa in Australia compared with other continents (Fiorini et al., 2023; Kerbs et al., 2017). By contrast, predominantly African *D.* sect. *Margaritaria* appear to be morphologically more diverse, and this clade also seems to be of hybrid origin. However, because little is known about the region, it is likely that the area is undersampled. Overall it should be noted that the morphological characters discussed here are notoriously highly variable even intraspecifically in Amaranthaceae (Sukhorukov & Zhang, 2013), which may be a consequence of a more relaxed selection on floral traits due to their predominantly anemophilous nature.

### Biogeography and landscape evolution

Statistical models suggest that *Dysphania* reached Australia either from North America, subsequently spreading to Eurasia and Africa, or from Eurasia, with later dispersal to Africa. While it cannot be ruled out completely, we here advocate for the second interpretation. American species appear to have a basic chromosome number of 8 and occur possibly exclusively as polyploids, whereas Eurasian and Australian taxa are likely exclusively diploid, with a basic chromosome number of 9 (Uotila et al., 2021). Furthermore, this dispersal route has been observed multiple times in other taxa that sympatrically occur with *Dysphania*: *Atriplex* (Žerdoner Čalasan et al., 2022), Camphorosmeae (Berasategui et al., 2024), *Triodia* (Anderson et al., 2019) and potentially *Gunniopsis* (Klak et al., 2017) and *Ptilotus* (Hammer et al., 2021) as well. The available time divergence estimation data suggest a similar timeframe in these lineages, which further solidifies this interpretation. In all available cases, long-distance dispersal has been proposed as the mechanism by which some of the ancestors of contemporary lineages reached Australia, given the lack of a fossil record to prove otherwise. The last common ancestor of Australian *Dysphania* probably arrived in the coastal regions of the contemporary Western Desert during the second half of the Miocene (Figures 3 and 4). There is ample evidence to suggest that this area was already markedly drier than the rest of the continent at that time, with most of the land covered by sclerophyll vegetation and some gallery forests, and with palaeodrainage systems becoming more irregular (Kershaw et al., 1994; Macphail et al., 1994; van de Graaff et al., 1977), characteristics of riverine desert habitats. These are characterised by desert drainage systems with occasional channel flows and infrequent inundation of broad floodplains with sheet flooding occurrences across the alluvial plains during periods of abundant rainfall (McDonald, 2020) and clearly represent the ancestral habitat of the Australian *Dysphania* (Figure 5).

Eastward expansion into the interior followed periods of increasing aridity and more frequent burning events (Figure 4). These events substantially transformed the Australian landscape, replacing rainforests with less dense sclerophyll forests, woodlands and chenopod shrublands (Martin, 2006). Similarly, the subsequent northward expansion of *Dysphania*, dated to the Pliocene, corresponds with palaeobotanical evidence indicating a reduction in tree cover and the gradual development of desert chenopod shrublands and grasslands (Martin & McMinn, 1994), despite the presence of a monsoonal system (Hesse, 2010; Figure 4). During the Pliocene, megalakes began to fragment into smaller basins (Cohen et al., 2011; Sniderman et al., 2016) as well and with increased evapotranspiration and reduced water influx, they subsequently started to shrink to their current size (Fitzsimmons et al., 2013). Repeated cycles of expansion and contraction of lake waterbodies throughout the Pleistocene may have initiated, or at least influenced, the speciation of *Dysphania* species in clade 2, which mainly occur along coastlines and the shores of inland lakes, a scenario that has also been proposed for other biota occurring in or along the shorelines of inland lakes (Bradford et al., 2013; López-López et al., 2016; Rahman et al., 2025; Žerdoner Čalasan et al., 2026)

While not restricted exclusively to these areas, most species of clade 1 are found in riverine deserts. Although riverine deserts originated in the western part of the continent, progressive aridification throughout the Late Neogene and Quaternary periods led to their eastward shift and disappearance from much of their former range (Mabbutt, 1988). Based on our habitat reconstruction (Figure 5), we postulate that the ancestor of contemporary Australian *Dysphania* at least to some extent retained its habitat and subsequently expanded eastward together with the riverine deserts. These extensive riverine systems are now established near the foothills of the Great Australian Divide. These landscapes remain relatively dry, however, they depend heavily on water from annual monsoonal or alpine flows (McDonald, 2020). Consequently, they form transitional zones where watercourses support more vegetation than the surrounding arid areas. The ecological similarities between riverine deserts and mesic vegetation across the Great Australian Divide in south-eastern and eastern Australia may explain why this clade includes species that have successfully colonised these regions, including climatically similar areas such as Tasmania and New Zealand. This ecological flexibility, which enabled them to survive in areas with unpredictable water supply and to thrive in disturbed habitats, may explain why this is the only clade of native Australian species with distributions beyond their native continent. Although the distribution of contemporary species does not support the littoral connection hypothesis, this signal may be obscured by the inland expansion of riverine deserts and the subsequent migration of the ancestor, which tracked this habitat rather than changing its ecology, as observed, for example, in some Camphorosmeae (Berasategui et al., 2024).

### Drivers of diversification

Our analysis suggests that *Suckleya* remained diploid, while a polyploidisation event gave rise to *Dysphania* (Supplementary File S6). An additional polyploidisation event seems to have occurred only in the last common ancestor of the American species. This finding is supported by chromosome counts, with *Suckleya* (Bassett & Crompton, 1970), Eurasian *D. botrys*, African/Arabian *D. schraderiana* and Australian *D. pumilio* and *D. carinata* being diploid (Grozeva & Cvetanova, 2013; Kawatani & Ohno, 1962) whereas the two American species *D. ambrosioides* and *D. anthelmintica* being polyploid (Grozeva & Cvetanova, 2013; Kawatani & Ohno, 1950), respectively. As we did not sample the entire taxon comprehensively, some polyploidisation events may not be captured by our taxon sampling and therefore cannot be inferred for lineages outside of Australia. However, polyploidisation evidently does not seem to play an important role in speciation of Australian *Dysphania* (Supplementary File S6).

Given the lack of clear geographic barriers in the relatively flat Australian landscape, allopatric speciation seems rather unlikely, and diverse soil and salinity profiles probably favours ecological speciation as a more plausible driver of diversification, as suggested for other Australian taxa (Anderson et al., 2016; Crisp & Cook, 2013; Pepper et al., 2013). This may also apply to many *Dysphania* species, which, despite occupying similar habitats, differ in their ecological niches (Supplementary Files S9 and S10). *Dysphania cristata* and *D. melanocarpa* are notoriously difficult to distinguish morphologically, differing mainly in tepal shape. While *D. cristata* occurs exclusively in riverine desert habitats, its spherical fruiting perianth may trap air pockets within multicellular hairs, maximizing buoyancy for hydrochory across flooded areas. In contrast, *D. melanocarpa* appears to occupy a slightly broader ecological niche and produces a bluntly stellate fruiting perianth, which may facilitate anemochoric dispersal in drier habitats where *D. cristata* does not occur. *Dysphania plantaginella* and *D. sphaerosperma* both occur along salt lakes and coastlines, with the former having a much wider distribution, whereas the latter is more restricted, typically associated only with gypseous or calcareous soils. *Dysphania rhadinostachya* seems better adapted to drier conditions with its erect habit, occurring mostly on skeletal soils on rocky outcrops, whereas *D. kalpari*, though morphologically similar, is prostrate and as such appears to be better adapted to occasionally flooded areas (Flowers & Colmer, 2015).

Literature shows that extensive hybridisation occurs frequently within the youngest clade 1 (Wilson, 1984), which radiated around 1.5 mya, whereas no comparable evidence exists outside this clade. Their recent divergence may explain why these species still hybridise upon secondary contact, whereas many other sympatric species that diverged earlier no longer do so. All *Dysphania* species are annuals or short-lived perennials, which makes them effective drought avoiders. Furthermore, the Australian *Dysphania* sect. *Dysphania* is characterised by markedly smaller habitus and seeds than other taxa (Uotila et al., 2021), possibly in response to the scarcity of reliable water and nutrient resources in arid environments. Furthermore, although evidence is limited, at least some species appear to exhibit heterospermy, producing seeds with different germination strategies on the same plant (Vandelook et al., 2021; Žerdoner Čalasan & Kadereit, 2023). This phenomenon is common in desert plants or species that occupy highly disturbed environments where favourable germination conditions are unpredictable (Baskin et al., 2014; Venable, 1985).

## Conclusion

Our results reveal a close evolutionary relationship between Sub-Saharan African and Australian desert ephemerals and indicate that the evolutionary history of Australian *Dysphania* was shaped by an ancestral reticulation event. The last common ancestor of *Dysphania* reached the coastal regions of northwestern Australia during the Miocene, occupying riverine desert habitats and initially migrating eastward alongside the expanding habitats, before potentially undergoing ecological speciation. Four major Australian clades subsequently diversified in response to distinct landscape phases, from Miocene riverine deserts to Pleistocene salt lake mosaics, where soil salinity gradients, flood frequency, and microhabitat partitioning putatively drove divergence without evident polyploidy or geographic isolation.

## Supporting information

Supplementary_File_S1_Voucher_information

Supplementary_File_S2_Australian_subregions

Supplementary_File_S3_ASTRAL_unreduced

Supplementary_File_S4_phyparts

Supplementary_File_S5_QS_analysis

Supplementary_File_S6_ASTRAL_orthogroup_mapping

Supplementary_File_S7_time_div_est_unreduced

Supplementary_File_S8_BioGeoBears_stats

Supplementary_File_S9_ENM_map_clade_1

Supplementary_File_S9_ENM_map_clade_2

Supplementary_File_S9_ENM_map_clade_3

Supplementary_File_S9_ENM_map_clade_4

Supplementary_File_S10_niche_overlap_results

## Supporting Information

**Supplementary File S1:** Voucher list of the taxa analysed in this study.

**Supplementary File S2:** Summed phytogeographical subregions according to Ebach et al. (2015), González-Orozco et al. (2014), and New Zealand. Areas are coded as follows: A, Hampton, Southwest Interzone, and Southwestern; B, Western Desert; C, Eastern Desert; D, Adelaide, Eyre Peninsula, and Nullarbor; E, Southeastern, Tasmania, Victoria; F, Eastern Queensland; G, Central Queensland; H, Great Sandy Desert Interzone and the Central Desert; I, New Zealand. The dashed line indicates that the distance between Australia and New Zealand is not proportional to that between Australia and Tasmania.

**Supplementary File S3:** ASTRAL-based phylogenomic tree of *Dysphania*, including additional samples across the Caryophyllales. Numbers on the branches indicate local posterior probabilities.

**Supplementary File S4:** Concordance analysis of Australian *Dysphania* inferred using PhyParts, plotted on the coalescent-based species tree. Colors indicate: blue, concordant topology; red, discordant topology; green, main alternative discordant topology; light grey, missing data; dark grey, uninformative data.

**Supplementary File S5:** Concordance analysis of Australian *Dysphania* inferred using Quartet Sampling, mapped onto the coalescent-based species tree. At each node, the three numbers indicate: quartet concordance (QC), quartet differential (QD), and quartet informativeness (QI).

**Supplementary File S6:** Orthogroup mapping results (in bold) showing potential gene duplications based on the subclade orthogroup tree topology method, plotted on the coalescent-based species tree.

**Supplementary File S7:** Dated phylogeny of *Dysphania*, including additional samples across the Caryophyllales. Numbers in square brackets at the nodes indicate the 95% HPD of estimated node ages.

**Supplementary File S8:** BioGeoBEARS model statistics for the spatial reconstruction of Australian *Dysphania* at both the global and continental scales.

**Supplementary File S9:** Ecological niche modelling maps of closely related Australian *Dysphania* species, with predicted habitat suitability indicated by a heatmap.

**Supplementary File S10:** Statistical comparison of ecological niche overlap among closely related Australian *Dysphania* species.

## Acknowledgements

We thank the curator of the CHR herbarium, I. Schönberger, for providing samples of the New Zealand endemic *Dysphania pusilla*, and A. Höwener (LMU München) for supervising the laboratory work conducted by the second author. We thank H. Štorchová (Institute of Experimental Botany, Czech Academy of Sciences) for providing pre-publication access to the transcriptomic data of *C. ficifolium* and P. Uotila (Finnish Museum of Natural History) for his comments on an earlier version of the manuscript.

## Author contributions

A.Z.C.: conceptualisation, methodology, formal analysis, figures, writing – original draft, data curation; F.S.: acquisition of data, writing – review and editing; K.K.: methodology, writing – original draft; B.M.: writing – review and editing; G.K.: conceptualisation, funding acquisition, writing – original draft.

## Data availability

The raw target enrichment data generated for this study are available under the NCBI BioProject PRJNAxxxxxxx. All code used in this study is available at https://bitbucket.org/dfmoralesb/target_enrichment_orthology.

