## Supplementary figures and images for "Tracing Sticky Trails: The Historical Biogeography of Australia’s Glandular Goose-foots (*Dysphania*, Chenopodioideae, Amaranthaceae)"

### Supplementary_File_S2_Australian_subregions

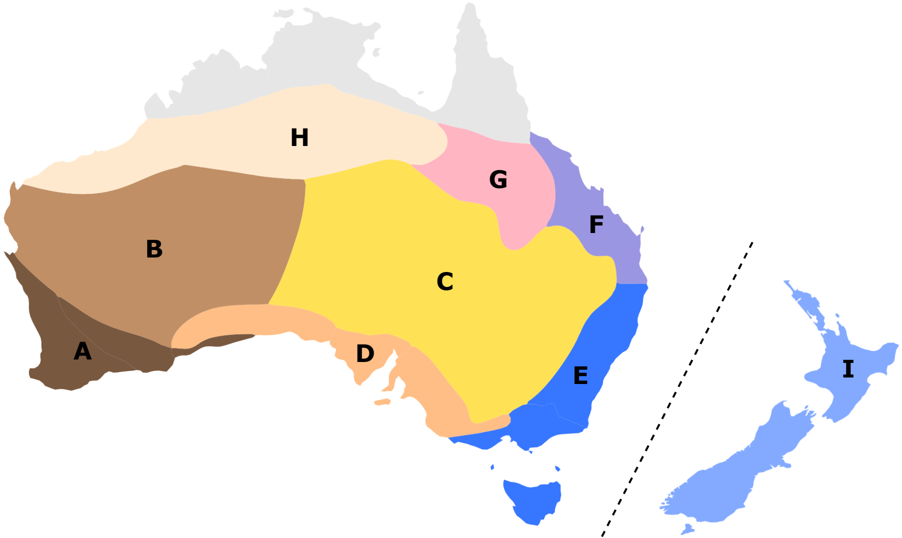

### Supplementary_File_S3_ASTRAL_unreduced

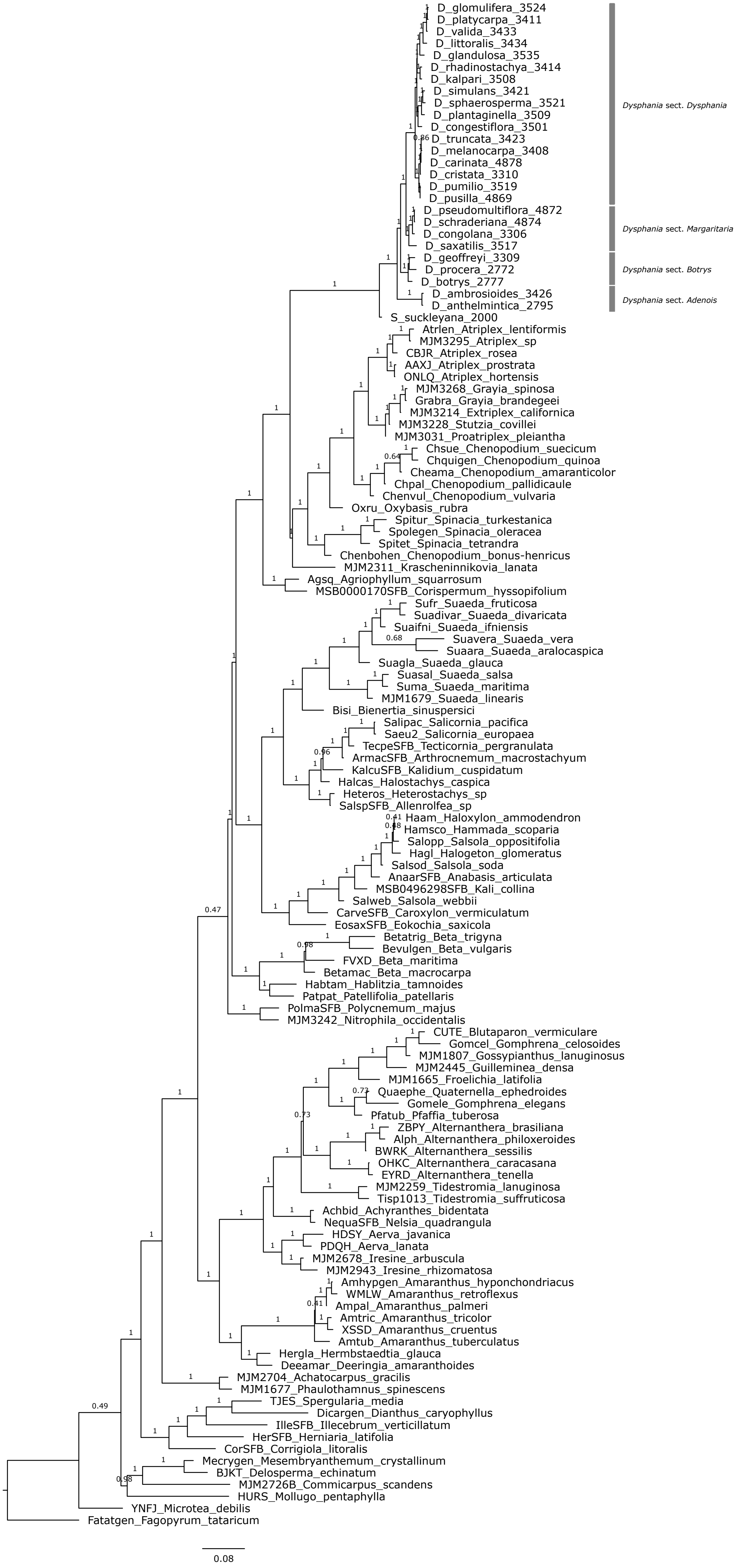

### Supplementary_File_S4_phyparts

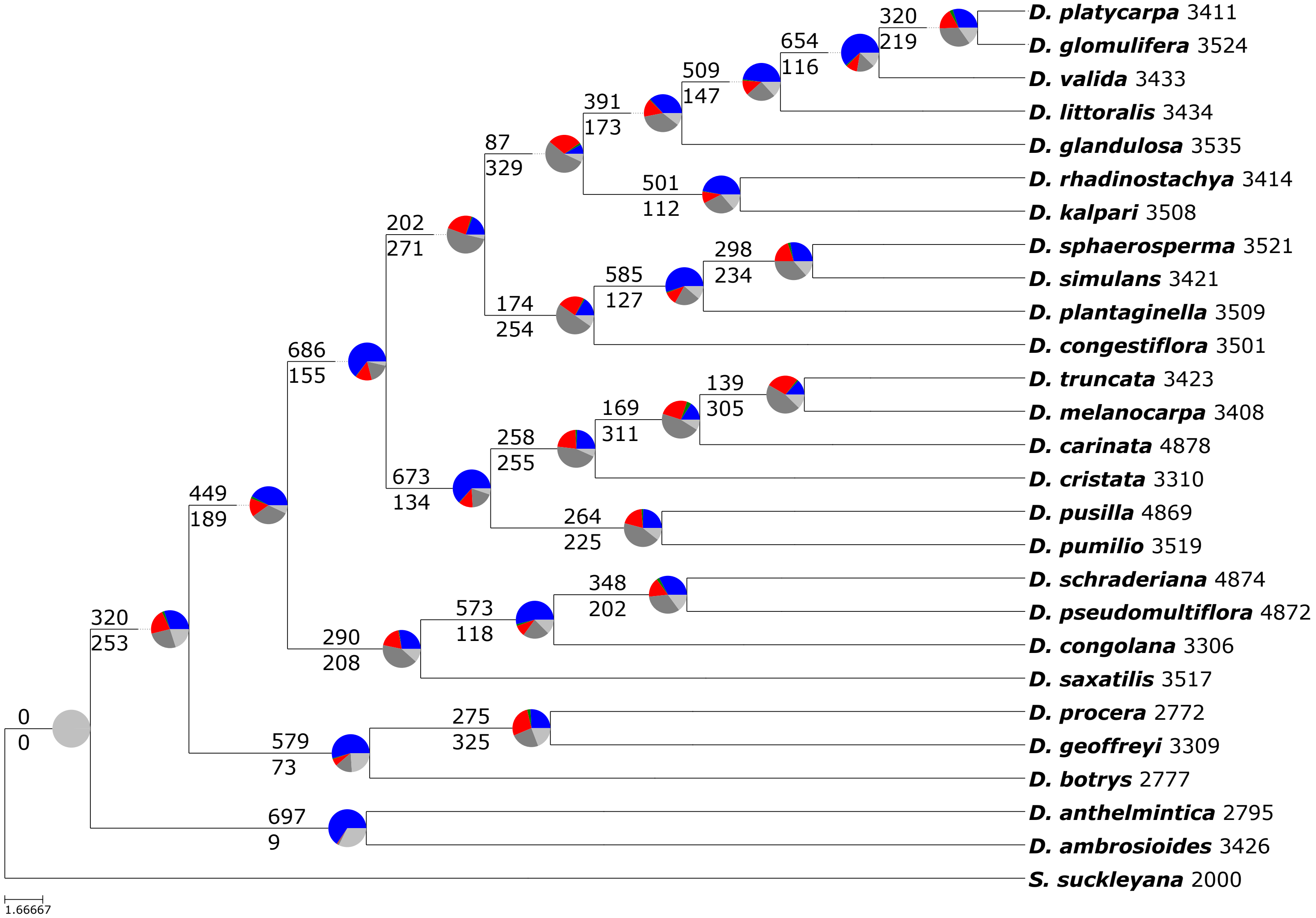

### Supplementary_File_S5_QS_analysis

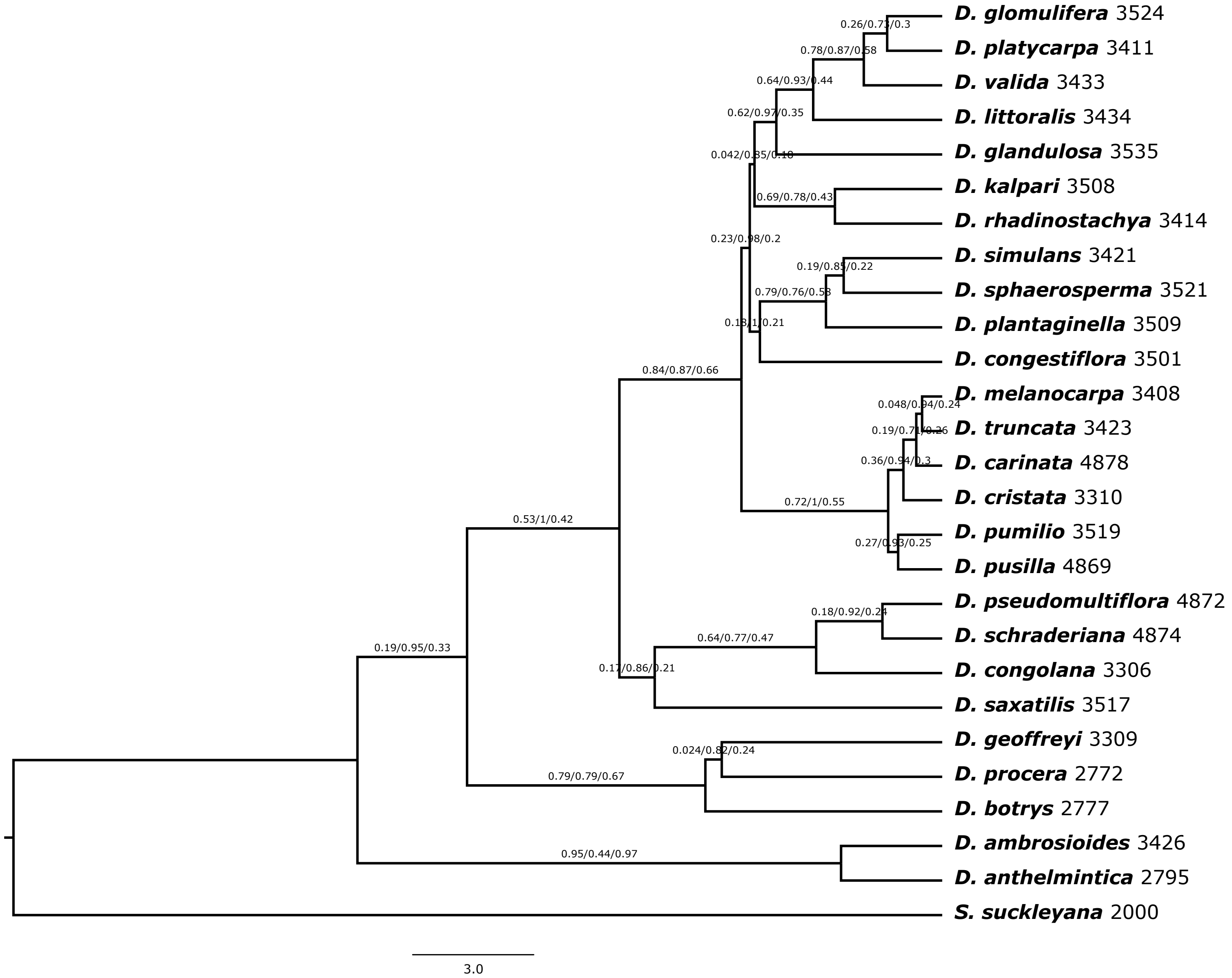

### Supplementary_File_S6_ASTRAL_orthogroup_mapping

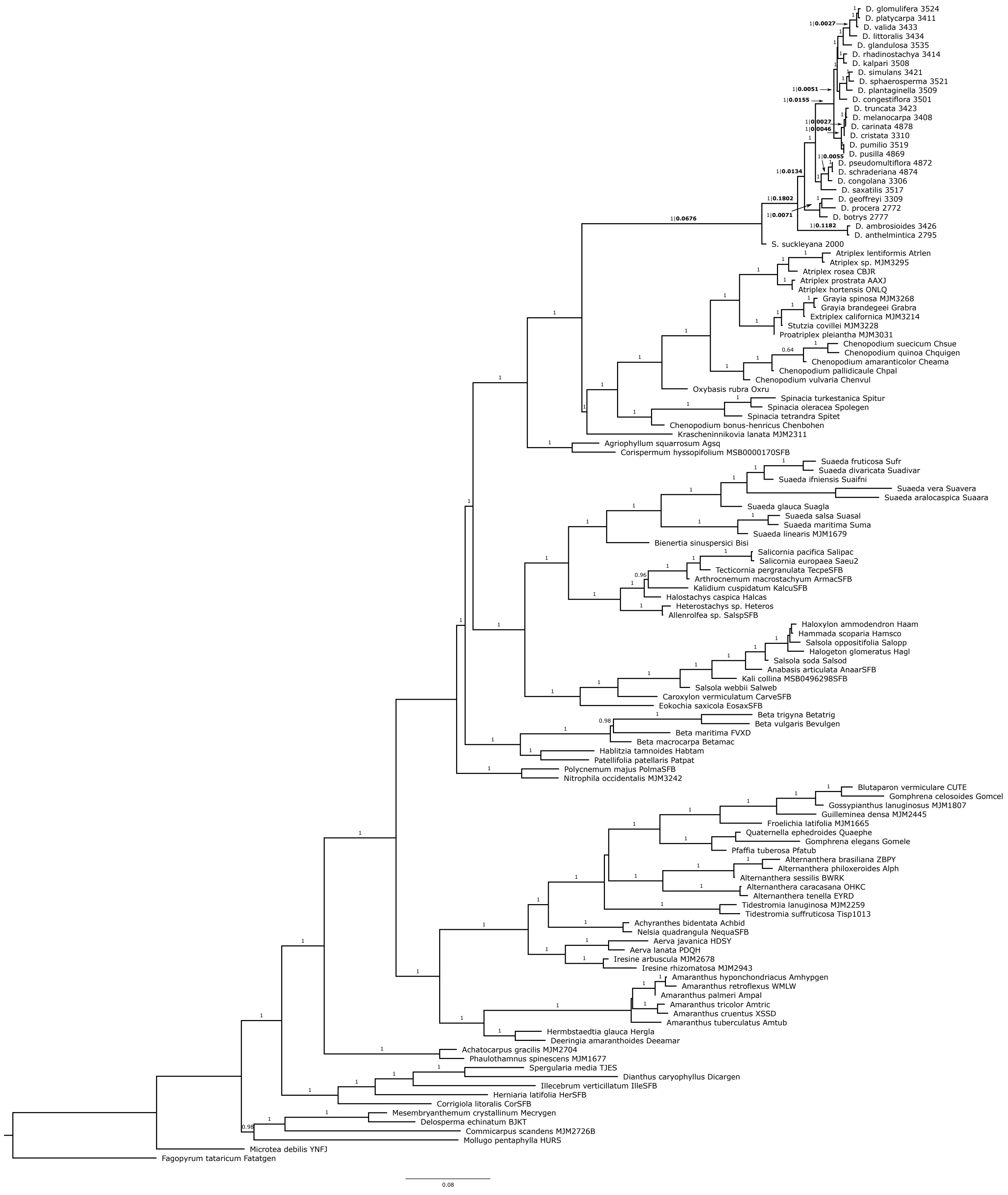

### Supplementary_File_S7_time_div_est_unreduced

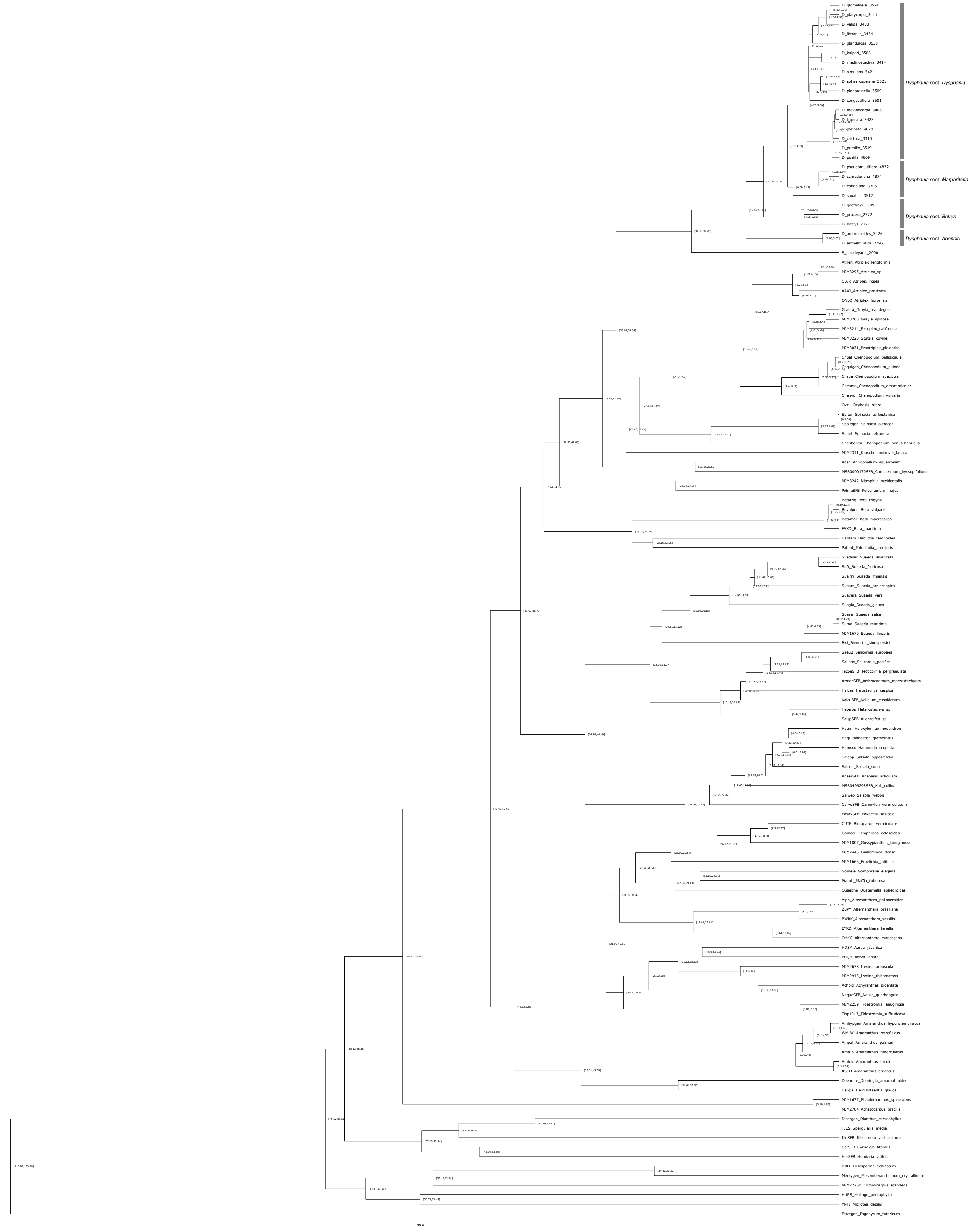

### Supplementary_File_S9_ENM_map_clade_1

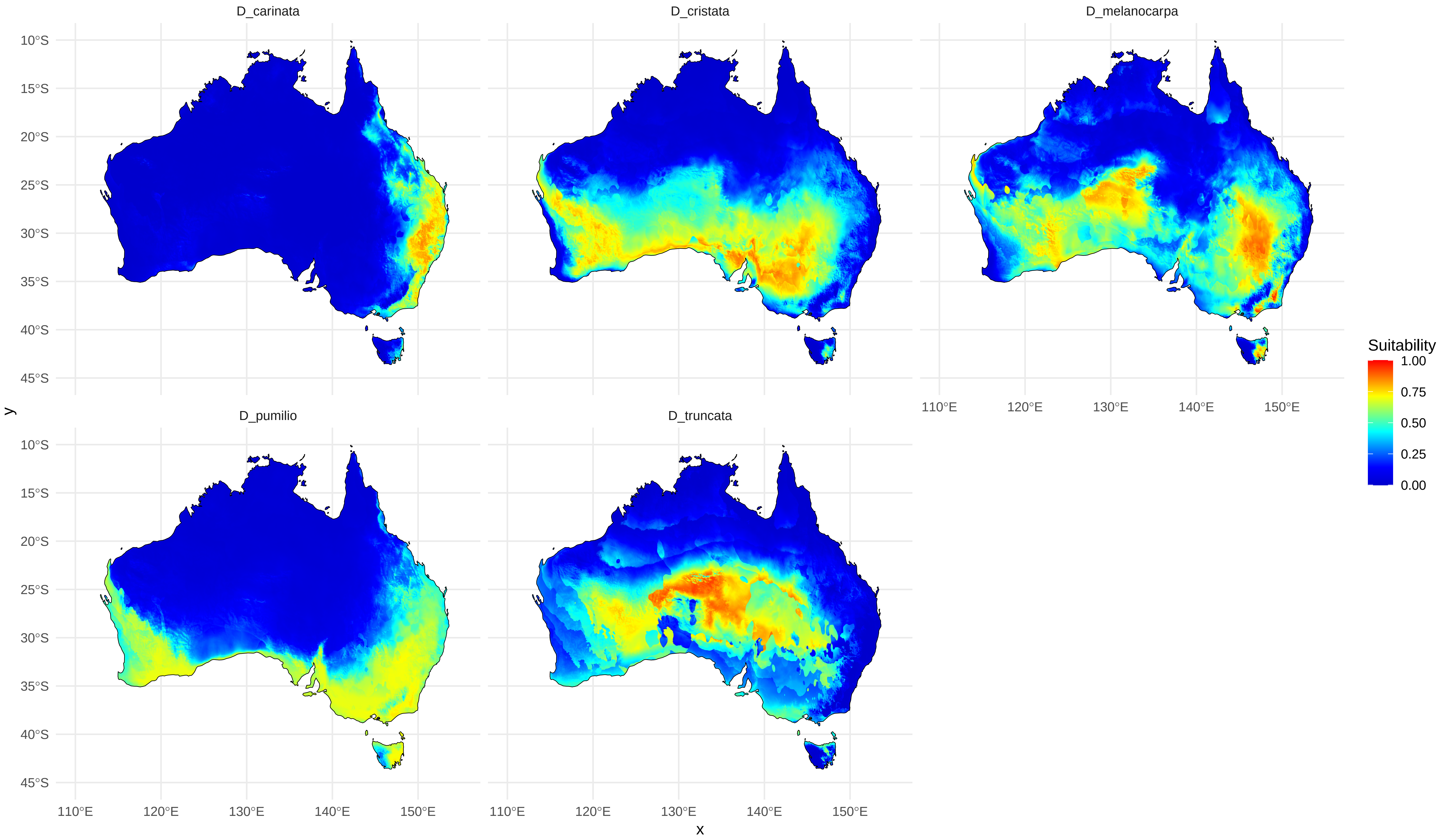

### Supplementary_File_S9_ENM_map_clade_2

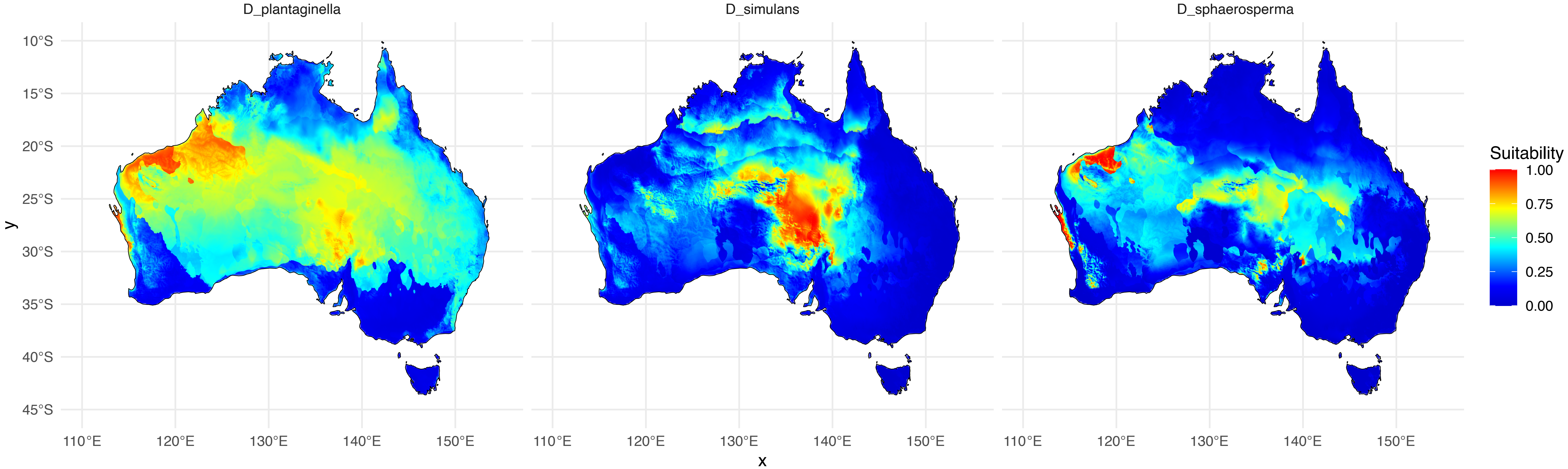

### Supplementary_File_S9_ENM_map_clade_3

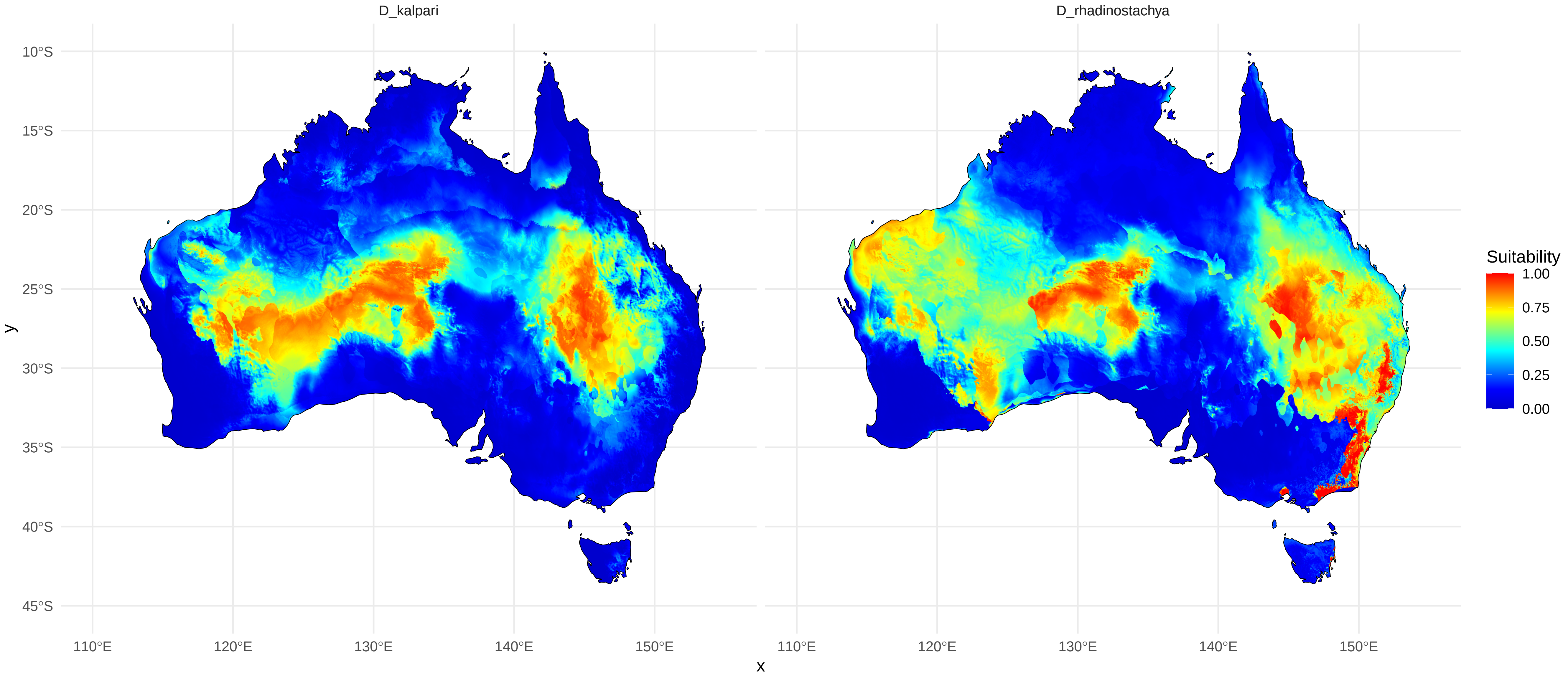

### Supplementary_File_S9_ENM_map_clade_4

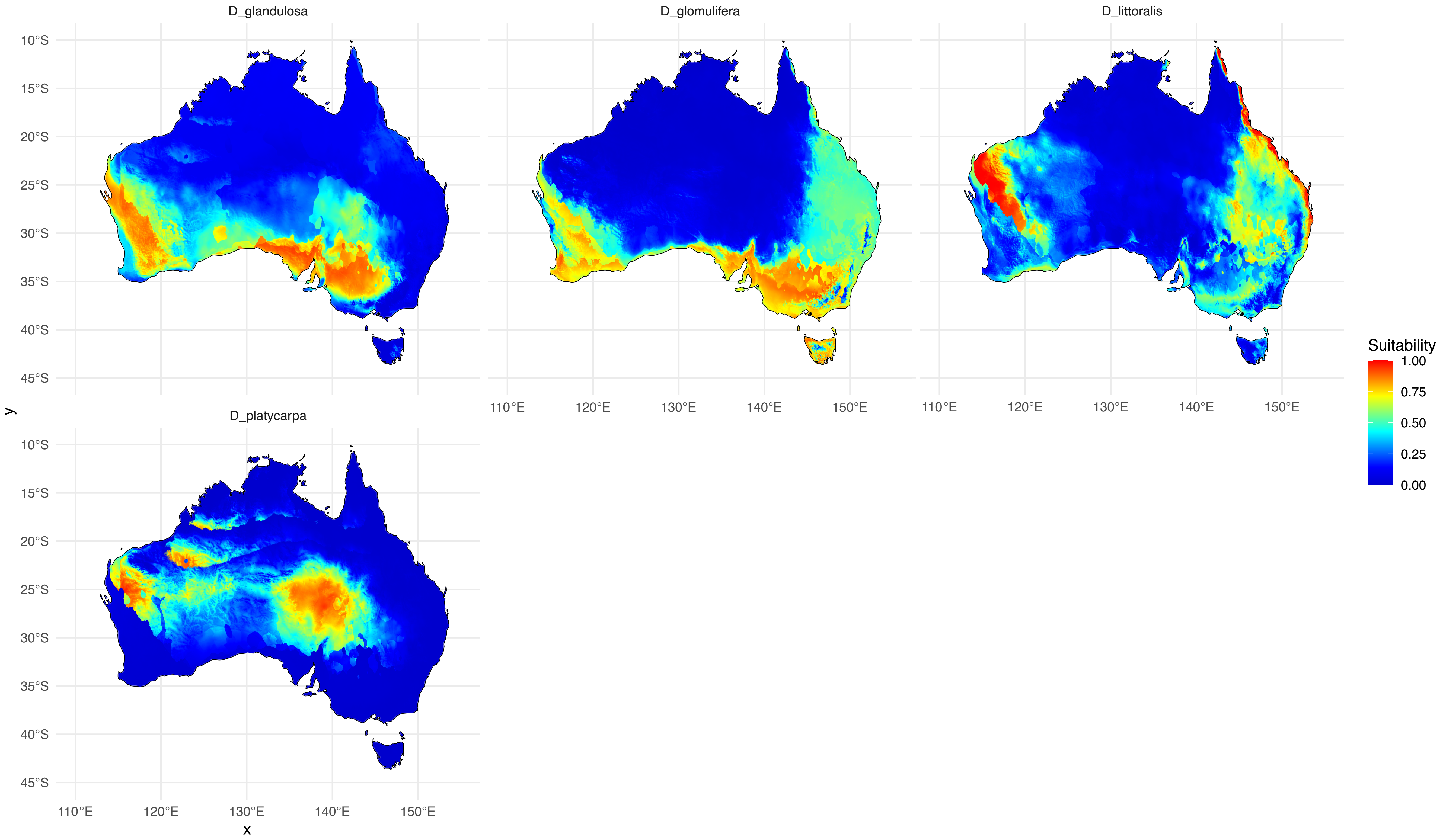
